# Overlapping and distinct roles of CDPK family members in the pre-erythrocytic cycle of the rodent malaria parasite, *Plasmodium berghei*

**DOI:** 10.1101/797886

**Authors:** K. Govindasamy, P. Bhanot

## Abstract

Invasion of, development within, and exit from hepatocytes by *Plasmodium* is essential for the parasite to establish the malaria-causing erythrocytic cycle. Identification of signaling pathways that operate during the pre-erythrocytic cycle provides insights into a critical stage of infection and potential targets for chemoprevention of disease. Calcium Dependent Protein Kinases (CDPK) represent a kinase family that is present in *Plasmodium* but absent in mammals. We demonstrate that *P. berghei* homologs of CDPK1, CDPK4 and CDPK5 play overlapping but distinct roles in sporozoite invasion and parasite egress from hepatocytes. All three kinases are expressed in sporozoites. All three are required for optimal motility of sporozoites and consequently their invasion of hepatocytes. Increased cGMP compensates for the functional loss of CDPK1 and CDPK5 during sporozoite invasion but cannot overcome CDPK4’s loss. CDPK1 and CDPK5 expression is downregulated after sporozoite invasion. CDPK5 reappears in a subset of late stage liver stages and is present in all merosomes. Chemical inhibition of CDPK4 and depletion of CDPK5 in liver stages suggests that these kinases play a role in the formation and/or release of merosomes from mature liver stages. Furthermore, depletion of CDPK5 in merosomes significantly delays a merosome-initiated erythrocytic cycle without affecting the infectivity of hepatic merozoites. These data suggest that CDPK5 is required for the release of hepatic merozoites from merosomes. Our work provides the first evidence that sporozoite invasion requires CDPK1 and CDPK5 and that the release of hepatic merozoites is a regulated process.

**Significance:** The malaria-parasite *Plasmodium* begins its mammalian cycle by infecting hepatocytes in the liver. A single parasite differentiates into tens of thousands of hepatic merozoites which exit the host cell in vesicles called merosomes. Hepatic merozoites initiate the first round of erythrocytic infection that leads to disease symptoms. We show that optimal invasion of liver cells by *Plasmodium* requires the action of three closely-related parasite kinases, CDPK1, 4 and 5. Loss of any of the three enzymes in the parasite significantly reduces infection of liver cells. CDPK5 is also required for the release of hepatic merozoites from merosomes and therefore for initiating the erythrocytic cycle. A better understanding of how these kinases function could lead to drugs that prevent malaria.

## Introduction

Parasite genomes are generally small in size with a great deal of functional optimization. The presence of gene families suggests that in some cases, structurally related proteins play redundant or complementary functions. One such gene family in the malaria-causing parasite, *Plasmodium* encodes Calcium Dependent Protein kinases (CDPK) that in addition to Apicomplexan parasites is found only in plants, protists, green alga and oomycetes (1). CDPKs are serine-threonine kinases that are activated by the direct binding of Ca^2+^ to EF-hand domains in their regulatory region. The ability to bind Ca^2+^ directly enables CDPKs to act both as sensors and effectors of intracellular Ca^2+^.

The human-infective species, *P. falciparum* encodes at least 7 members (2). The rodent-infective species, *P. berghei* encodes orthologs for all members except for *P. falciparum* CDPK2. The function of different CDPKs have been extensively examined in the asexual cycle in erythrocytes and during parasite transmission to and development within mosquitoes. These studies have revealed distinct and overlapping roles for different CDPK members during merozoite invasion and egress, gametogenesis and ookinete invasion. In the asexual cycle, CDPK5 is required for merozoite egress (3, 4), and CDPK1 and CDPK4 play non-essential roles in merozoite invasion (5, 6). Although invasion by *P. berghei* merozoites is reduced when CDPK4 is lost together with decreased function of cGMP dependent protein kinase (PKG) but the loss of CDPK4 alone does not affect asexual growth of *P. berghei* or of *P. falciparum* (6). Similarly, in the background of simultaneous decrease in PKG and CDPK4 functions, CDPK1’s loss decreases merozoite invasion (6, 7) but its individual loss is tolerated in the asexual cycle (6, 8, 9). CDPK2, CDPK3 and CDPK6 do not appear to play significant roles in the asexual cycle (10–13). During the parasite’s sexual cycle in the mosquito midgut, CDPK1, CDPK2 and CDPK4 are essential for male gametogenesis with CDPK4 regulating at least 3 distinct cell cycle events (13–18). Ookinete motility requires CDPK3 (10, 11) as it regulates the secretion of adhesins required for motility. CDPK1 and CDPK4 also contribute to ookinete motility: simultaneous loss of both reduced ookinete speed *in vitro* (6). These studies of different CDPK family members in the mosquito transmissive stages has provided biological validation for targeting *P. falciparum* CDPK1 and CDPK4 to chemically block parasite transmission from mammalian host to mosquitoes.

Sporozoites and liver stages express several CDPKs (8, 12, 19) but, in contrast to our knowledge in asexual and sexual cycles, little is known of CDPKs’ functions during the pre-erythrocytic cycle. CDPK4 and CDPK6 function in invasion by *P. berghei* sporozoites since their loss reduces the percentage of sporozoites that enter hepatocytes (12, 19) while deletion of CDPK1 in sporozoites did not reveal a significant defect in parasite invasion of or egress from hepatocytes (8). These results suggested that other CDPKs could play a compensatory and complementary role during sporozoite invasion and parasite exit from hepatocytes.

Here we examine the role of CDPK1, CDPK4 and CDPK5 in sporozoite motility, traversal through, invasion of and egress from infected hepatocytes, using conditional protein depletion in the rodent model, *P. berghei*. *P. berghei* offers the distinct advantage of an experimentally accessible pre-erythrocytic cycle. We demonstrate that in *P. berghei* sporozoites, CDPK1 (PBANKA_0314200), CDPK4 (PBANKA_0615200) and CDPK5 (PBANKA_1351500) are individually required for motility. The loss of each kinase decreases sporozoite motility and consequently significantly reduces their invasion of hepatocytes. CDPK5 is required for parasite egress from hepatocytes and for release of hepatic merozoites from merosomes. Our study is the first demonstration of CDPK1’s and CDPK5’s role in sporozoite invasion. Furthermore, it provides evidence that release of hepatic merozoites from merosomes is a regulated process that requires CDPK5.

## Results

### CDPK1, CDPK4 and CDPK5 have dynamic expression patterns during the pre-erythrocytic cycle

Since CDPK1, 4, 5 have essential roles in the asexual and sexual cycles, genetic analyses of their functions in pre-erythrocytic stages requires conditional mutagenesis. We adapted a method for conditional protein degradation previously described in *P. berghei* asexual stages and ookinetes. Here, a target protein is tagged with an Auxin-induced degron (AID) in Ostir1-expressing transgenic *P. berghei* (20). The tag targets the protein for rapid proteasomal degradation upon the addition of the plant hormone, auxin (Indole-3-acetic acid (IAA). We constructed parasite lines, CDPK4-aid-HA and CDPK5-aid-HA, in which either CDPK4 or CDPK5 were fused with an AID domain and tagged with an HA_2x_ epitope (Supplementary Fig. 1A, B). These two parasite lines and a previously reported CDPK1-aid-HA line (20) were transmitted to mosquitoes for recovery of sporozoites from salivary glands.

Microscopic examination of midguts revealed fewer oocysts in mosquitoes infected with CDPK1-aid-HA or CDPK4-aid-HA parasites compared to mosquitoes infected with an isogenic control (data not shown). Oocyst numbers in mosquitoes infected with CDPK5-aid-HA were equivalent to control (data not shown). Since genomic deletions of CDPK1 and CDPK4 abolish male gametogenesis (15, 16), decreased numbers of oocysts in CDPK1-aid-HA and CDPK4-aid-HA lines suggest that the AID-HA_2x_ domain either interferes with the activities, expression or localization of CDPK1 and CDPK4 and therefore, partially impairs gametogenesis in these lines. Consistent with reduced midgut infection in CDPK1-aid-HA and CDPK4-aid-HA and normal midgut infection in CDPK5-aid-HA, there was a significant decrease in the average number of salivary gland sporozoites in CDPK1-aid-HA or CDPK4-aid-HA lines but the decrease in ones infected with CDPK5-aid-HA was not significant (Supplementary Fig. 1C). Despite reduced numbers of CDPK1-aid-HA and CDPK5-aid-HA sporozoites, we obtained sufficient parasites to conduct our analyses. Due to the low numbers of CDPK4-aid-HA sporozoites, their analysis was more limited.

We began functional interrogation of CDPKs in the pre-erythrocytic cycle by examining their temporal and spatial protein expression in sporozoites and liver stages. We previously reported the presence of CDPK4 in sporozoites and intracellular liver stage parasites formed 24-65 h post infection (p.i.) of HepG2 cells (19). Here we demonstrate that CDPK1 and CDPK5 are also present in sporozoites (Fig. 1) and are distributed throughout the cytoplasm. In liver stages, expression of the three kinases is dynamic. CDPK1 and CDPK5 were not detected in early liver stages 24-48 h p.i (Fig. 1). At 65 h p.i., when liver stages are close to exiting the hepatocyte, CDPK5 was detected in a subset of liver stages where it sometimes partially co-localized with *P. berghei* PKG (PbPKG) at the periphery of the infected cell (Fig. 1B). CDPK5 expression correlated with developmental maturity since, in the same culture, liver stages that express it were significantly larger in size compared to those that did not have detectable CDPK5 (Fig. 1C). CDPK5’s appearance in mature liver stages suggested its expression may coincide temporally with parasite egress from the infected hepatocyte. Therefore, we examined its expression in merosomes and detached cells released at 65-67h p.i. CDPK5 was detectable in all merosomes/ detached cells and localized to the merosomes periphery, most likely at the membrane (Fig. 2). PbPKG was also present in at the merosome periphery but the two kinases did not display significant co-localization.

**Figure 1.**
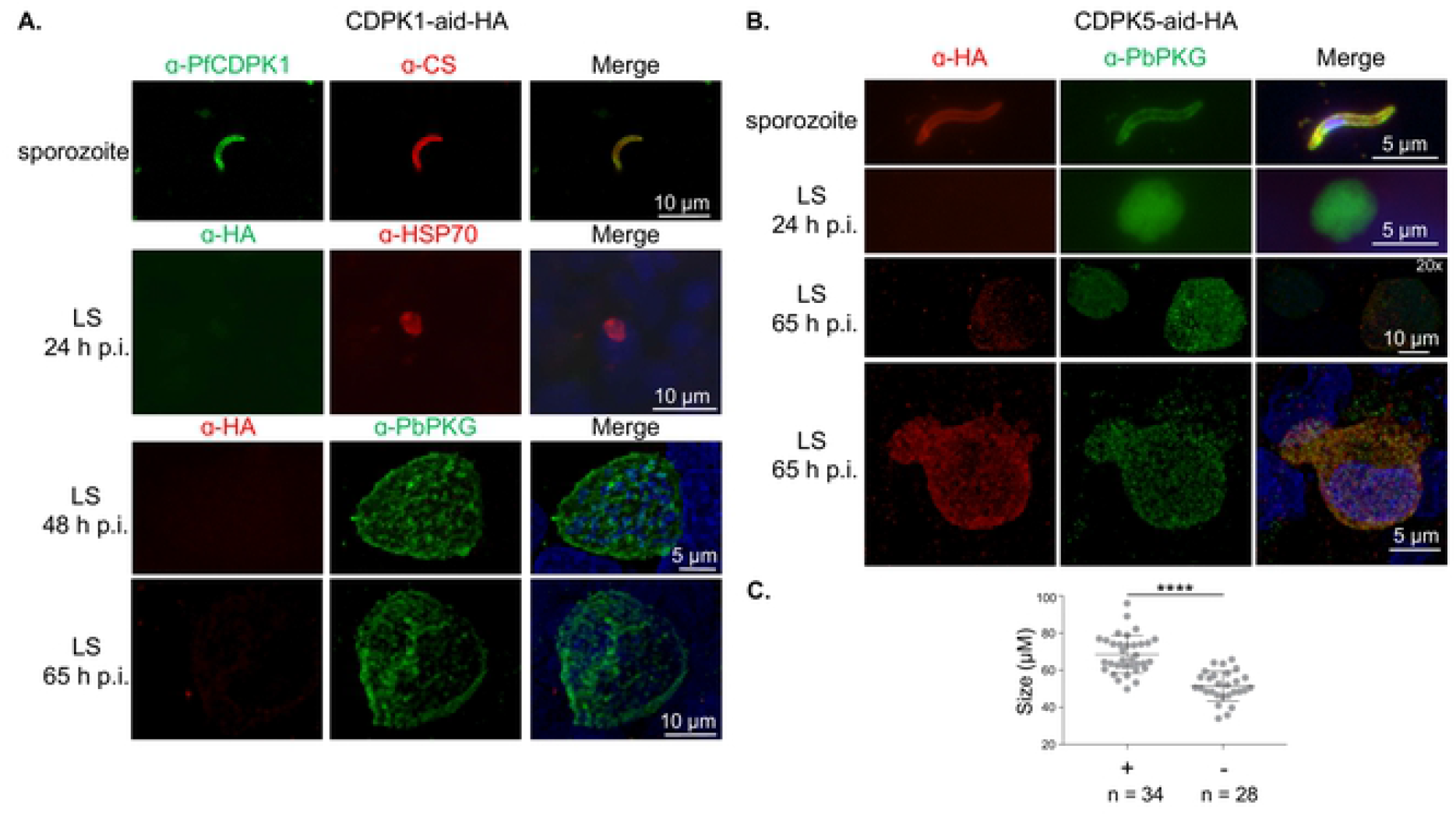
Spatial and temporal localization of CDPK1 and CDPK5 during the pre-erythrocytic cycle of *P. berghei*. **A)** Representative images of CDPK1 localization in pre-erythrocytic stages of CDPK1-aid-HA parasites was determined using anti-HA or anti-P. *falciparum* CDPK1 (PfCDPK1) antibodies. Sporozoites were co-stained with CS and liver stages with either Heat Shock Protein 70 (HSP70) or cGMP dependent protein kinase (PbPKG). CDPK1 was present only in sporozoites **B)** Representative images of CDPK5 localization in pre-erythrocytic stages of CDPK5-aid-HA parasites determined using an anti-HA antibody. CDPK5 was present in all sporozoites and in a subset of mature liver stages (65 h p.i.). Its expression was undetectable in early liver stages (24 h p.i.). **C)** Diameter of individual CDPK5-aid-HA liver stages (μM ± SEM), that either displayed staining with anti-HA antibody at 65h p.i. (+) or not (-) was determined. Data were analyzed using an unpaired *t*-test, **** *P* value < 0.0001.

**Figure 2.**
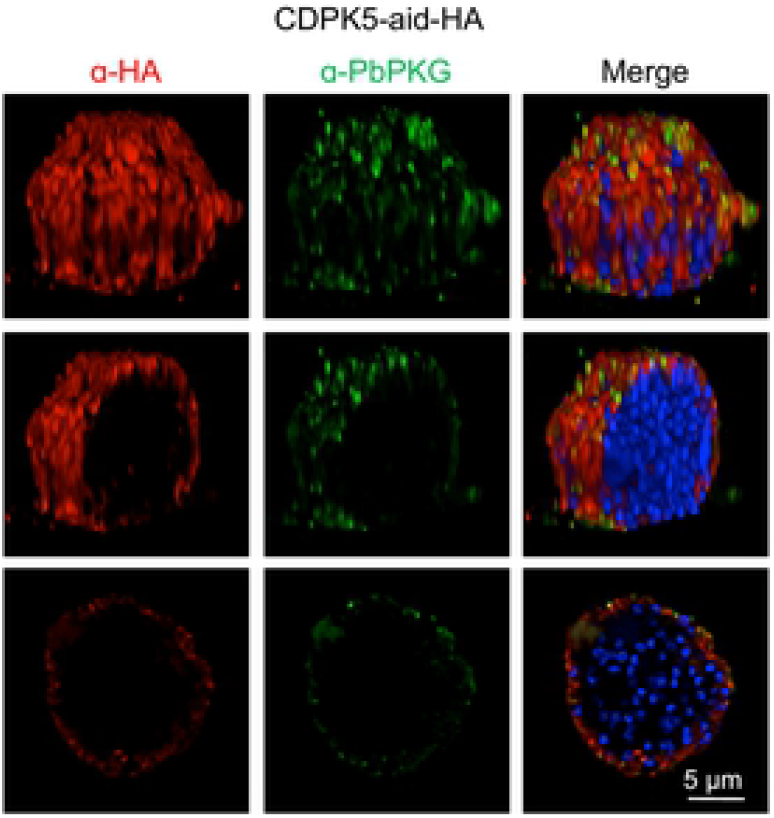
CDPK5 and PKG are present in the periphery of merosomes. Representative deconvolved images and optical sections of immunostained CDPK5-aid-HA merosomes.

We have previously shown that PbPKG is required for merosomes formation or release (19, 21). To obtain a better understanding of the subcellular localization of PbPKG in developing liver stages, we colocalized it with Exp1, a resident protein of the parasitophorous vacuole membrane, HSP70, a cytoplasmic protein and MSP1, a resident protein of the parasite plasma membrane (Fig. 3). PbPKG colocalized with Exp1 but not HSP70 or MSP1. Taken together, these data suggest that in developing liver stages, PbPKG associates with the vacuole membrane. After membrane breakdown in mature liver stages, PbPKG becomes associated with the hepatocyte membrane and is in close proximity to CDPK5. During merosomes formation, the two proteins are associated with the hepatocyte membrane that envelops hepatic merozoites as they are released from the infected cell in the form of merosomes.

**Figure 3.**
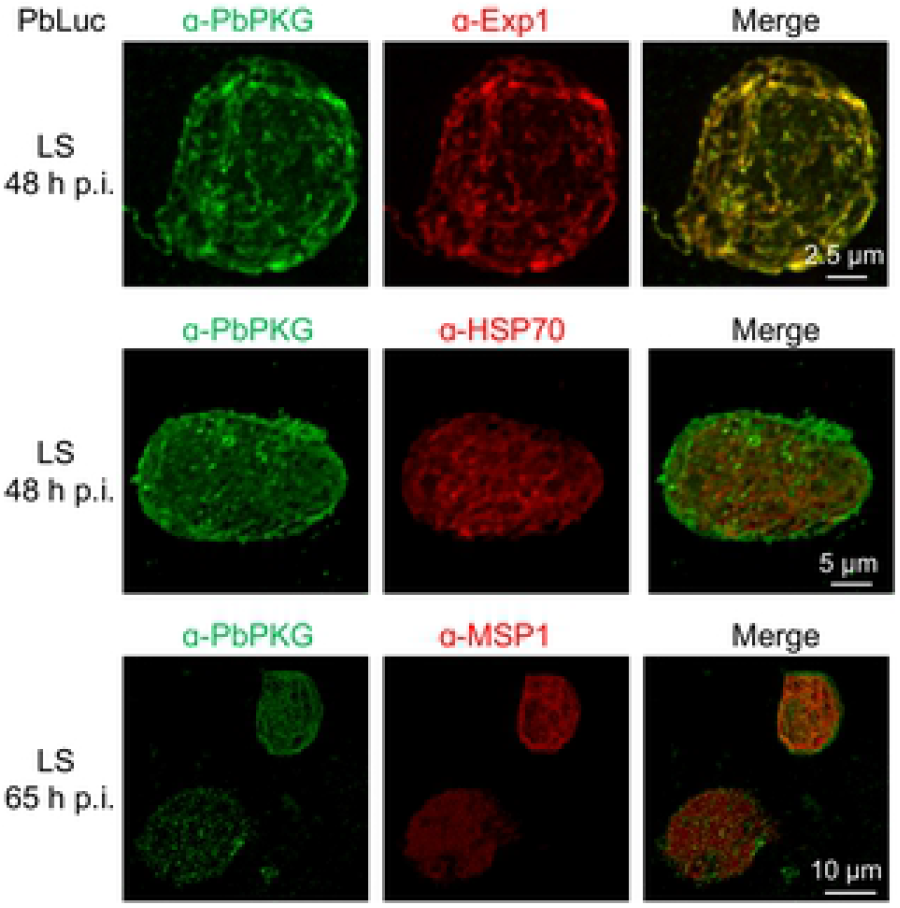
PbPKG is associated with the parasitophorous vacuole membrane of liver stages. Representative images of PbPKG’s spatial expression in liver stages of *P. berghei* parasites expressing luciferase (PbLuc). Exp1 is a resident protein of the parasitophorous vacuole membrane, HSP70 is a cytoplasmic protein and Merozoite Surface Protein (MSP1) is a marker of the parasite plasma membrane.

### Validation of AID-mediated conditional protein degradation in pre-erythrocytic stages

The unique expression profiles of CDPK1, 4 and 5 in pre-erythrocytic stages suggested some overlapping but also distinct roles. We tested the function of the three kinases in the pre-erythrocytic cycle using conditional protein depletion. We tested the efficiency of IAA-mediated protein degradation in CDPK1-aid and CDPK5-aid sporozoites. In both cases, Western blot analysis of protein lysates demonstrated an almost complete loss of AID-tagged proteins (Fig. 4A). Since CDPK5-aid-HA protein migrated faster than its predicted molecular weight, we confirmed its loss in IAA-treated sporozoites using immunofluorescence assays (Supplementary Fig. 1D). In addition, we confirmed its loss in released merosomes treated with IAA (Fig. 4B).

**Figure 4.**
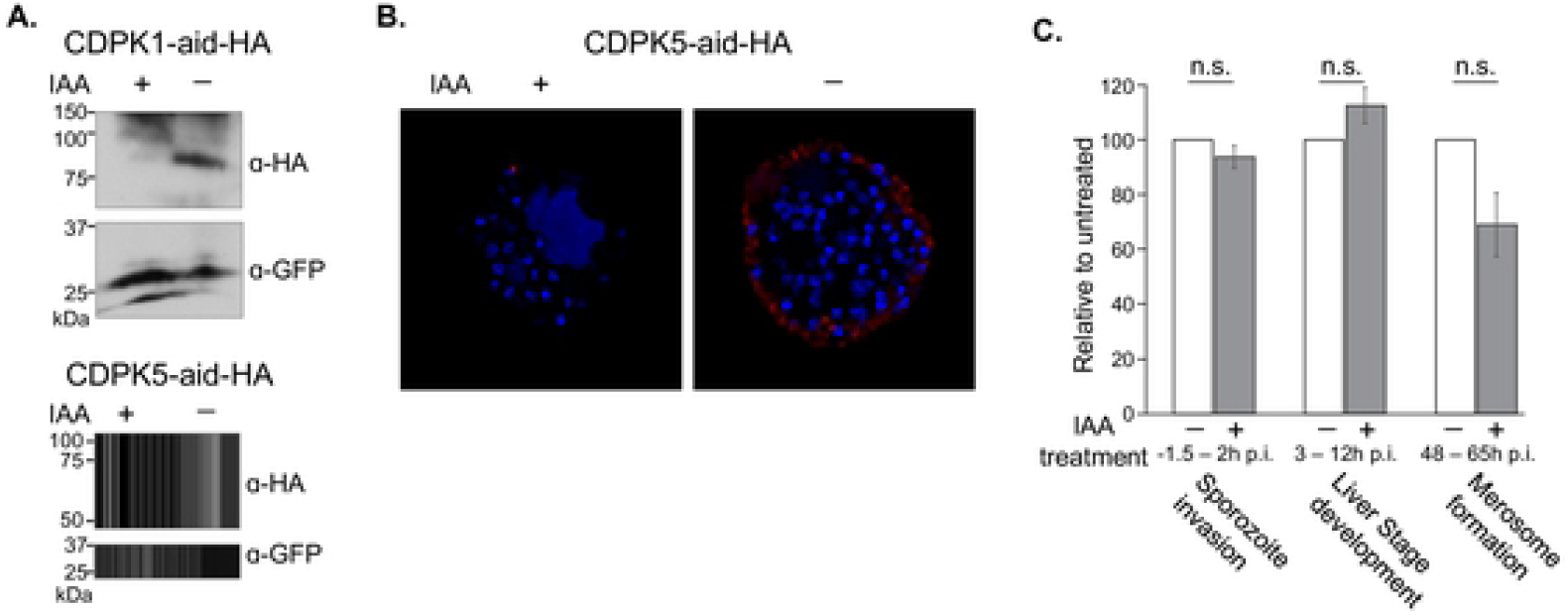
Conditional degradation of AID-tagged proteins is efficient in pre-erythrocytic stages. **A)** Western blot analysis of protein lysates from CDPK1-aid-HA and CDPK5-aid-HA sporozoites treated with IAA or vehicle. CDPK1-aid-HA and CDPK5-aid-HA were detected using an anti-HA antibody. GFP was used as a loading control. CDPK1-aid-HA and CDPK5-aid-HA have predicted molecular weights of approximately 90 kDa. **B)** Depletion of CDPK5-aid-HA protein in merosomes was examined using immunofluorescence. Representative images of IAA- or vehicle-treated merosomes stained with an anti-HA antibody. Parasite nuclei were visualized using DAPI. **C)** Effects of IAA treatment on Ostir1 pre-erythrocytic stages were examined by quantifying sporozoite invasion of HepG2 cells, the number of liver stages detected at 48 h p.i. and the number of merosomes released into the media at 66-70h p.i., in the presence or absence of IAA. Results shown are average of 2-4 experiments with 3 technical replicates (± SEM), normalized to vehicle-treated samples for each assay. Data were analyzed using an unpaired *t*-test, not significant (n.s.) *P* value > 0.05.

Next, we examined if conditions required for protein degradation of AID-tagged proteins have non-specific effects on parasites. We tested IAA’s effect on motility, invasion, infectivity and egress of isogenic control sporozoites that lack AID-tagged proteins, Ostir1 (20). Motility was quantified by live imaging sporozoites after treating with IAA or vehicle (Movie 1-2). The percentage of Ostir1 sporozoites that moved in complete circles was similar in both conditions - 36% of vehicle-treated (n = 175) and 33% of IAA-treated (n =165). Sporozoite invasion was determined by quantifying the percentage of IAA- and vehicle-treated sporozoites that enter HepG2 cells within 90 min. Their intracellular development was assessed by quantifying the number of liver stages present in HepG2 cells after addition of IAA or vehicle to sporozoite-infected HepG2 cultures 2-14 p.i. (Fig. 4C). There was no significant effect of IAA treatment on sporozoite invasion or on liver stage development as the fraction of sporozoites that entered cells and the number of liver stages was similar in sporozoites treated with IAA or vehicle (Fig. 4C). Effect of IAA on parasite egress from hepatocytes was examined by quantifying the number of merosomes present in media of control-infected HepG2 cells treated with IAA 48-65 h p.i. IAA treatment did not significantly affect merosome release, demonstrating that it does not affect egress of control parasites. We conclude that IAA treatment specifically targets aid-HA proteins and does not have a deleterious effect on sporozoites. Therefore, IAA-mediated conditional degradation of proteins is a powerful tool for studying protein function in *Plasmodium’s* pre-erythrocytic cycle.

### CDPK1, CDPK4 and CDPK5 are required for sporozoite motility

*In vitro*, sporozoites display a variety of movement patterns – moving in continuous circles (gliding), waving with one end attached to the substrate, attached with no movement or a combination of the previous three (22). Motility is accompanied by dynamic changes in intracellular Ca^2+^ (23). Ca^2+^ flux regulates secretion of adhesins onto the sporozoite surface that mediate attachment to the substrate, and turnover of adhesion sites between the sporozoite and substrate during movement (23, 24). To examine the role of CDPK1, CDPK4 and CDPK5 as Ca^2+^ effectors during motility, we examined the effect of their loss on sporozoite motility.

Depletion of CDPK1, CDPK4 or CDPK5 had no significant effect on sporozoite attachment to the substrate but sporozoites no longer demonstrated circular movement (Movie 3-8, Fig. 5A). Instead, they stayed attached to the substrate with either one pole (‘waving’) or both poles attached (22). These results suggest that the formation and efficient turnover of attachment sites between the sporozoite and the substrate, and consequently sporozoite motility, requires a threshold of Ca^2+^ signaling that is reached through the combinatorial function of each of these kinases. Each kinase can partially but not completely compensate for the loss of another.

**Figure 5.**
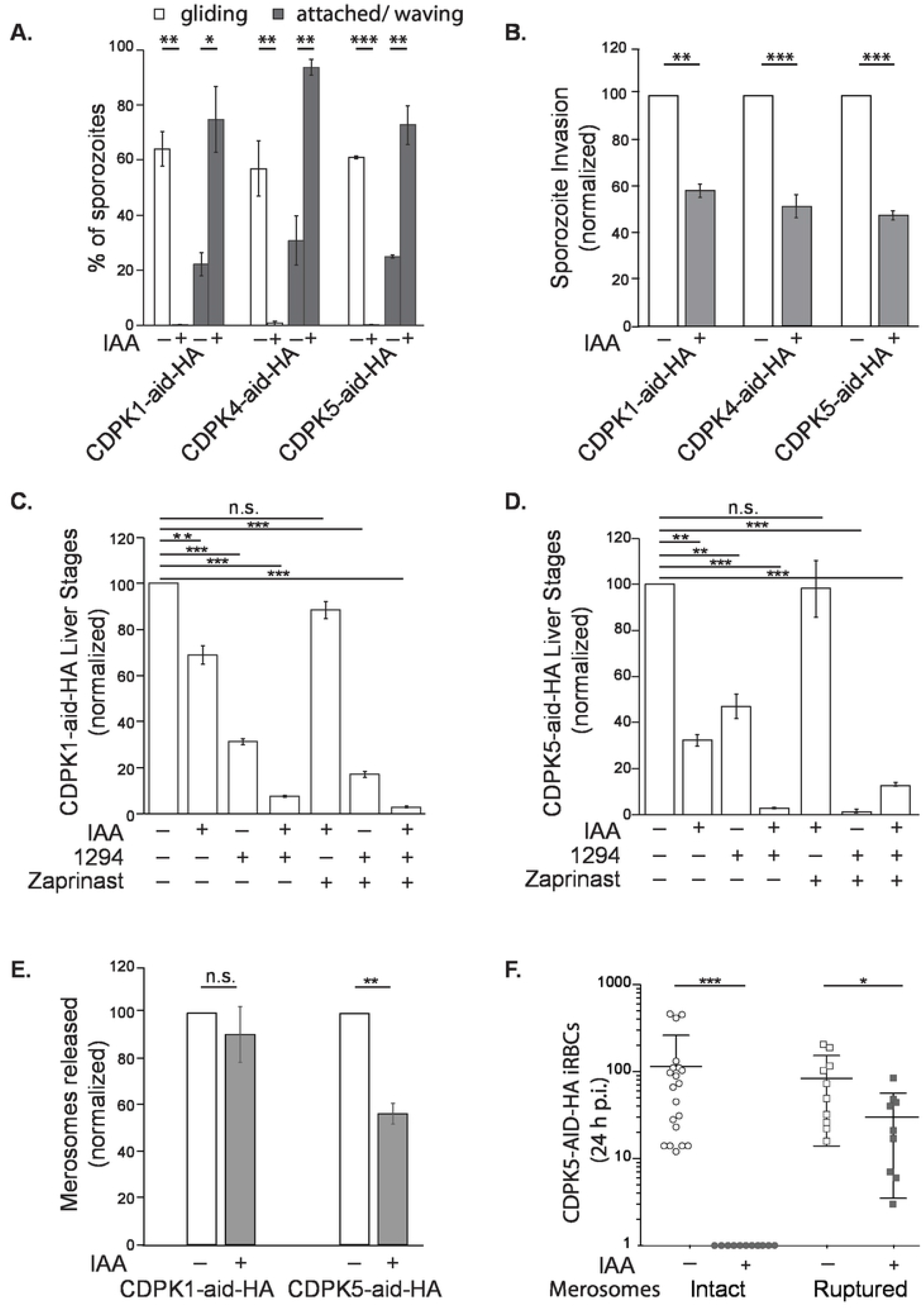
CDPKs have partially overlapping but distinct roles during the pre-erythrocytic cycle. **A)** Gliding motility of sporozoites was quantified as the percentage of sporozoites that engage in circular motion (± SEM). The experiment was performed 3-4 times for each parasite line. **B)** Sporozoite invasion was examined by determining the percentage of sporozoites that become intracellular within 90 min of addition to HepG2 cells. Results shown are average of 3-4 experiments with 3-4 technical replicates (± SEM), normalized to vehicle-treated samples. **C)** Interaction between CDPK1, CDPK4 and PKG in sporozoites was studied by determining the number of liver stages formed by sporozoites in which CDPK1 and CDPK4 functions are inhibited, when PKG is activated. Results shown are average of 3 experiments with 3 technical replicates (± SEM), normalized to vehicle-treated samples. **D)** Interaction between CDPK5, CDPK4 and PKG in sporozoites was studied by determining the number of liver stages formed by sporozoites in which CDPK5 and CDPK4 functions are inhibited, in the presence of increased cGMP. Results shown are average of 3 experiments with 3 technical replicates (± SEM), normalized to vehicle-treated samples for each assay. **E)** Merosome formation and release was determined by quantifying the number of merosomes present in media of infected cells at 65-67 h p.i. Results shown are average of 2 experiments with CDPK1-aid-HA and 3 experiments with CDPK5-aid-HA, with 3 technical replicates (± SEM), normalized to vehicle-treated samples. **F)** Infectivity of CDPK5-aid-HA merosomes and hepatic merozoites was quantified by FACS detection of GFP-positive events in 100,000 events at 24 h p.i. (iRBCs ± SEM). The experiment was performed 4 times with intact merosomes (5 mice/each group/experiment), and 2 times with ruptured merosomes (4-5 mice/each group/experiment). Data were analyzed using an unpaired *t*-test or one-way ANOVA, Dunnett’s multiple comparisons test, non-significant (n.s.) *P* > 0.05, * *P* value < 0.05, ** *P* value < 0.005, *** *P* value < 0.0005.

*In vivo*, motility enables sporozoites to disseminate from the site of bite in the skin by traversing through cell layers, entering a blood vessel and invading a hepatocyte in the liver. Therefore, we examined if decreased motility that results from CDPK1, 4 or 5 depletion has any functional effect on cell traversal by sporozoites and their eventual invasion of hepatocytes. Depletion of CDPK1 or CDPK5 did not decrease cell traversal by sporozoites (Supplementary Fig. 1E). We were unable to test cell traversal in CDPK4-aid-HA sporozoites due to insufficient sporozoites. Invasion of HepG2 cells decreased by approximately 50% in all IAA-treated sporozoites (Fig. 5B). These data demonstrate that CDPK1, CDPK4 and CDPK5 are required for sporozoite invasion of hepatocytes and they have partially overlapping functions in the process. Optimal invasion by sporozoites requires all three kinases. The ability of sporozoites to invade cells despite depletion for CDPK1 or CDPK4 or CDPK5 and an apparent complete loss of circular motion *in vitro* likely suggests that motility on glass coverslips is likely an incomplete representation of sporozoite motility on extracellular matrix. The extracellular matrix might provide a greater number or variety of adhesins using which sporozoites can achieve sufficient motility.

Decreased invasion by CDPK1-depleted sporozoites is expected to diminish the number of liver stages formed by these sporozoites. However, previous studies did not find a significant difference in the number of liver stages formed by sporozoites in which the CDPK1 gene was deleted using stage-specific DNA excision (CDPK1 cKO) (8). To reconcile these seemingly contradictory results, we examined the possibility that CDPK1 protein is not significantly reduced in CDPK1 cKO sporozoites, using immunofluorescence assays (Supplementary Fig. 2). Protein was clearly detected in these sporozoites and its presence was entirely consistent with the normal infectivity of CDPK1 cKO sporozoites. In contrast, IAA-mediated CDPK1-aid-HA protein degradation is more efficient (Fig. 4A). Therefore, conditional degradation of CDPK1 uncovered its function in sporozoite motility and by extension, in host cell invasion for which motility is a pre-requisite.

To determine the effect of simultaneous loss of CDPK1 and CDPK4, we generated a parasite line in which both kinases are tagged with AID-HA (CDPK1-aid-HA/CDPK4-aid-HA) (data not shown). CDPK1-AID-HA/CDPK4-AID-HA parasites were viable in the erythrocytic cycle and were transmitted to mosquitoes through a blood meal but they failed to form any salivary gland sporozoites (data not shown). This negative interaction between modified alleles of CDPK1 and CDPK4 confirms previous reports of epistatic interactions between the kinases during gametogenesis (6).

As an alternative, we utilized a CDPK4-specific bumped kinase inhibitor compound 1294 (19, 25) to test the effect of simultaneously inhibiting CDPK4, and CDPK1 or CDPK5. Addition of 1294 (2 μM) to IAA-treated CDPK1-aid-HA and CDPK5-aid-HA sporozoites at the time of invasion significantly decreased the number of intracellular liver stages present at 48 h p.i., compared to liver stages formed by sporozoites treated with IAA or 1294 alone (Fig. 5C-D). We tested the possibility that 1294’s effect was partly through off-target inhibition of PbPKG, which has a small amino acid at the gatekeeper position. *P. berghei* sporozoites expressing PbPKG with a gatekeeper substitution (T_619_Q) remain sensitive to 1294 demonstrating that PbPKG is not a major target of the compound (Supplementary Fig. 3A). Together, these data suggest that CDPK1, CDPK4 and CDPK5 act together to provide optimal sporozoite invasion.

Since CDPKs act downstream of PbPKG during merozoite invasion of erythrocytes (CDPK4 and CDPK1 (6, 7) and egress (CDPK5 (4)), we tested if increasing cGMP levels, and consequently PbPKG signaling, during sporozoite invasion could overcome the loss of CDPK1, CDPK4 or CDPK5. Zaprinast, a phosphodiesterase inhibitor, reversed the effect of depleting either CDPK1 or CDPK5 individually (Fig. 5C-D). The number of liver stages formed by CDPK1-depleted or CDPK5-depleted sporozoites in the presence of zaprinast was indistinguishable from vehicle-treated sporozoites. However, zaprinast could not overcome simultaneous inhibition of CDPK4 and either CDPK1 or CDPK5 sporozoites (Fig 6C-D). These results suggest that PbPKG acts upstream of both CDPK1 and CDPK5 during sporozoite motility and that CDPK4 functions in a pathway that is at least in part independent of PbPKG.

### CDPK5 is required for parasite egress from hepatocytes and from merosomes

*Plasmodium’s* entry into the blood stream after exiting the liver is a two-step process. First, merosomes are extruded from the infected hepatocyte into the blood stream (26, 27). Second, merosomes are carried intact into the lung microvasculature where they disintegrate and release free hepatic merozoites that initiate the first round of erythrocytic infection (27). Egress from hepatocytes requires the activity of parasite PKG (27) and SUB1 (28) but there is little information on processes required for the release of hepatic merozoites from merosomes.

The presence of CDPK4 and CDPK5 in late liver stages suggests a possible function during parasite egress from hepatocytes. However, a conditional deletion of CDPK4 gene in sporozoites did not demonstrate a clear effect on the number of merosomes released in culture (19). We attempted to confirm these results using IAA-mediated depletion of CDPK4-aid protein in liver stages but the small number of CDPK4-aid-HA sporozoites made these assays unfeasible. As an alternative, we tested the effect of chemical inhibition of CDPK4 on merosome formation. Addition of 1294 to sporozoite-infected HepG2 cells at 48 – 66 h p.i. decreased merosomes released into media at 66 h p.i (Supplementary Fig. 3B). The different effects of genetic and chemical inhibition of CDPK4 suggest that parasites can adapt to genetic ablation of CDPK4 in sporozoites but cannot do so in response to chemical inhibition. Since chemical inhibition of kinase activity is rapid, it may not allow sufficient time for upregulation of compensatory pathways that can be upregulated in mutant parasites during their development from sporozoites into merosomes.

Depletion of CDPK5-aid in liver stages, starting 48 h p.i., also reduced the number of merosomes formed at 65h p.i. Similar treatment of CDPK1-aid-HA-infected HepG2 cells had no significant effect consistent with the lack of CDPK1 expression in liver stages (Fig. 5E). These results demonstrate that Ca^2+^ signaling through CDPK5 is required for parasite exit from hepatocytes. Since CDPK5 is detected in free merosomes (Fig. 2), we investigated its function in the release of hepatic merozoites. CDPK5-aid-HA merosomes treated with either IAA or vehicle prior to being injected intravenously into mice. In order to detect infection by hepatic merozoites, blood parasitemia was monitored at 24 h p.i. This time-period enables the initiation and completion of a single asexual cycle in *P. berghei*. None of the mice infected with CDPK5-depleted merosomes (n = 20) demonstrated parasitemia at 24 h p.i. while all mice infected with vehicle-treated merosomes were positive (Figure 5F). Mice infected with CDPK5-depleted merosomes became patent on day 3 p.i. (n = 20) and the growth rate of asexual parasites was similar in the two groups (Supplementary Fig 3C). We conclude that CDPK5’s loss in merosomes does not affect replication of asexual stages.

The longer pre-patent period and the lower parasitemia at 24h p.i. from CDPK5-depleted merosomes suggested that CDPK5 plays an important role in the release of hepatic merozoites from merosomes and/or in their invasion of erythrocytes. To distinguish between these two possibilities, we tested the effect of CDPK5 depletion of the infectivity of hepatic merozoites. We manually ruptured vehicle- or IAA-treated CDPK5-aid-HA merosomes prior to their injection into mice. In this case, all mice were patent at 24 p.i. and the difference in average parasitemias of the two groups of mice was much smaller (Figure 5F). Together, these results are consistent with a model in which CDPK5 functions in the rupture of merosome membrane and release of hepatic merozoites.

## Discussion

Initiation of the erythrocytic cycle by hepatic merozoites is possibly the least-understood step of the malaria infection cycle. Hepatic merozoites have a single opportunity to infect host cells and their release at the wrong time or place, for example in a non-vascular environment, would prevent or severely debilitate the launch of the erythrocytic cycle. While there is significant understanding at the molecular level of events leading to the release of merozoites from infected erythrocytes, almost nothing is known of how hepatic merozoites are released from merosomes. We provide strong evidence that release of hepatic merozoites from merosomes is a parasite-regulated process. The breakdown of the merosome membrane is likely a response to environmental cues that result in Ca^2+^ flux in merosomes, activation of CDPK5 and release of proteolytic enzymes.

Depletion of CDPK5 from merosomes does not impair the ability of hepatic merozoites to invade erythrocytes since manually-released hepatic merozoites from CDPK5-depleted merosomes initiate erythrocytic infection normally. In contrast, intact merosomes depleted for CDPK5 exhibit a significant delay in initiating erythrocytic infection as demonstrated by the significantly lower blood stage parasitemia in mice 24 h p.i.. It has been suggested that a lung-receptor mediated mechanism arrests merosomes in lungs where infection of erythrocytes by hepatic merozoites is facilitated by the low macrophage density and reduced shear forces in the pulmonary capillary bed from lower blood velocity (27). We hypothesize that interaction between merosomes and a tissue-specific receptor could trigger CDPK5-mediated release of hepatic merozoites. Another mechanism that could account for delayed patency of CDPK5-depleted merosomes is better clearance by the host immune system. Testing these models requires development of quantitative assays for release of hepatic merozoites from merosomes. In addition, it will be interesting to explore if CDPK5’s function during hepatic merozoite release requires CDPK5 to localize to the merosome membrane.

CDPK5 is also required for the formation of merosomes. The relatively modest effect on merosomes formation could suggest that CDPK5 primarily functions after merosomes have been released from the hepatocyte i.e. in the release of hepatic merozoites. Alternatively, CDPK4, which is also present in liver stages at 65 h p.i., could have a redundant role with CDPK5 in merosome formation. It is also possible that CDPK5 function in merosome formation is compensated by PbPKG. We cannot rule out the possibility that IAA-mediated depletion of CDPK5-aid-HA in intracellular and intravacuolar parasite stages, such as liver stages, is less efficient compared to its depletion in sporozoites or released merosomes. Incomplete depletion of CDPK5 in liver stages could provide sufficient protein for close-to-normal function in merosome formation.

The relative dearth of knowledge about the function of CDPKs in pre-erythrocytic stages can be attributed, in part, to the difficulties faced in functional analyses in pre-erythrocytic stages of kinases that have indispensable functions in the asexual and sexual cycles. Despite these challenges, it is important to understand functions of CDPKs in these stages because they constitute the first step of the mammalian infection, and the study of CDPKs’ in these stages could identify targets for malaria chemoprevention. In addition, it will provide a fuller view of the stage-specific and stage-transcending functions of different family members. The relative contribution of different family members in different parasite stages may be determined by the threshold of Ca^2+^ signaling required for different cellular processes, the sensitivity of specific kinases to Ca^2+^ level, their expression level and subcellular localization.

Our work reveals nuances of CDPK signaling in different parasite stages. First, our results demonstrate a new stage-specific role for CDPK5 - in sporozoite motility and consequently parasite invasion of hepatocytes. CDPK5 has not been implicated in RBC invasion by merozoites (4). In contrast to invasion, CDPK5’s role in parasite egress is stage-transcending as it is required for egress from both erythrocytes and hepatocytes. During the erythrocytic cycle, CDPK5 triggers micronemal secretion leading to merozoite egress from schizonts (4). How CDPK5 enables release of hepatic merozoites from merosomes remains to be investigated. Based on its localization to the merosomes membrane, it is tempting to speculate that CDPK5 could activate a spatially-restricted protease cascade close to the merosome surface.

Second, our results demonstrate that while different invasive stages may require the same set of CDPK kinases, the relative contribution of specific kinase members can vary at different stages or during specific processes. These differences may reflect the different host environments encountered by the different stages. We show that CDPK1, CDPK4 and CDPK5 have distinct roles in sporozoite motility and that these functions are only partially overlapping. Depletion of any of these kinases significantly reduces sporozoite motility. Since IAA-treated sporozoites attach to the surface, we speculate that their defective motility is likely due to dysregulation in turnover of adhesion sites that is required for the sporozoites to move forward (24). In contrast, in ookinetes, individual depletion or inhibition of CDPK1 and CDPK4 does not affect motility *in vitro* and only simultaneous loss significantly reduces ookinete speed (6). The implication is that each kinase can fully compensate for the other during ookinete traversal (6). As shown previously, ookinete motility and traversal through the midgut epithelium relies more heavily on CDPK3 (11).

We were surprised to find that, despite the loss of continuous movement, sporozoites depleted of CDPK1 or CDPK5 maintained the ability to traverse through cells albeit being significantly attenuated in invasion. Sporozoites traverse through cells by forming a transient vacuole without the formation of a tight junction between the sporozoite and hepatocyte membranes (29). Productive invasion requires the formation of a tight junction and sporozoites are contained within a parasitophorous vacuole. Since loss of CDPK1 and CDPK5 reduces only invasion, it implies that residual motility in depleted sporozoites is sufficient for traversal. We suggest that CDPK1 and CDPK5 could regulate the formation of the tight junction or another process specific to the form of cell entry that is accompanied by the formation of a permanent vacuole.

Optimal invasion by sporozoites requires a network of CDPKs – CDPK1, CDPK4 and CDPK5, as shown here, and CDPK6 as shown previously (12). We posit that, *in vivo*, reduced motility that results from inhibition or loss of CDPK1, CDPK4 or CDPK5 will severely impair cell entry by sporozoites, thereby attenuating hepatocyte infection. In contrast, during invasion by asexual stages, the loss of CDPK1 or CDPK4, individually or together, has no significant effect. The role of CDPK4 in erythrocytic invasion is seen only in the background of reduced PKG activity (6). CDPK1’s role in invasion is uncovered only in the background of simultaneous reduction in PKG and CDPK4 activities. It is possible that erythrocytic stages adapt to the loss of CDPK1 and CDPK4 by upregulating another CDPK member or PKG activity.

We have also determined that CDPK4 and CDPK5 play a role in merosome formation (Fig. 5E and Supplementary Fig. 3B). Although conditional deletion of the CDPK4 gene in sporozoites (CDPK4 cKO) did not result in a significant decrease in merosome formation (19), its chemical inhibition did. We consider it unlikely that 1294-mediated reduction in merosomes is due to off-target effects on PbPKG because 1294’s IC_50_ against PfPKG is a fold higher than its IC_50_ for PfCDPK4 and *P. berghei* sporozoites carrying a gatekeeper mutation in PbPKG remain sensitive to 1294 (Supplementary Fig 3A). The lack of merosome reduction in CDPK4 cKO parasites may be attributed to upregulation of compensatory pathways. To test these models, we wanted to determine the effect on merosome formation of rapid reduction in CDPK4 protein. We were unable to do so because CDPK4-aid-HA sporozoites are not produced in sufficient numbers for robust merosome assays. Further studies using alternative methods for depleting CDPK4 exclusively in liver stages and compounds with greater specificity for CDPK4 will address this issue.

## Materials and Methods

### Ethics statement

All animal work in this project was reviewed and approved by the Institutional Animal Care and Use Committee (IACUC) of Rutgers New Jersey Medical School, approval number TR201900067, following guidelines of the Animal Welfare Act, The Institute of Laboratory Animal Resources Guide for the Care and Use of Laboratory Animals, and Public Health Service Policy.

### Construction of AID-HA tagged parasite lines

The targeting plasmid for modifying CDPK4 with the aid-HA degron was constructed by amplifying a C-terminus fragment of CDPK4 using PCR primers AATTGGAGCTCCAC**CGCGG**CAAGTATTAAGTGGTATTACATATATG and TCATTCTAGT**CTCGAG**ATAGTTACATAGTTTTATTAACATGTCTC. The PCR product was cloned into the previously described AID-tagging plasmid expressing mCherry (pG364) (20), using SacII and XhoI. The targeting plasmid for modifying CDPK5 with the aid-HA degron was constructed by cloning a fragment amplified using PCR primers AATTGGAGCTCCA**CCGCGG**CATAGAGATTTAAAGCCAGAA and TCATTCTAGT**CTCGAG**AGATTGTCTTCCAGACATC, into a previously described AID-tagging plasmid expressing GFP (pG362) (20). The CDPK4-aid-HA targeting plasmid was linearized using BspEI and the CDPK5-aid-HA targeting plasmid was linearized using BclI. Linearized plasmids were transfected into the previously described OsTIR1-expressing parent line (20), using standard procedures (30). Transfected parasites were selected by pyrimethamine treatment and clonal lines were established through limited dilution. Modification of CDPK4 was confirmed using PCR primer pairs P1 (TGAAGTAGATGCAGCTAG) + P2 (GTTAAATGTGGGGTAAAAAA) and P3 (GTATTTACCCTGTCATACAT) + P4 (GATTAAGTTGGGTAACGC). Modification of CDPK5 was confirmed using PCR primer pairs P2 + P5 (CAAATGGATCATCCAAATATT) and (P3) + P6 (AAGGAATAGAAGGTAGAAATTG).

### Mosquito infections

*Anopheles stephensi* mosquitoes were fed on infected Swiss-Webster mice. Mosquitoes were maintained on 20% sucrose at 25°C at a relative humidity of 75-80%. Sporozoites were obtained by crushing salivary glands dissected on days 18–25 post-feeding. Sporozoites were counted in a hemocytometer.

### Immunofluorescence assays for protein detection in sporozoites, liver stages and merosomes

Primary antibodies used were: anti-HA (Biolegend, 1:200 −1:400), anti-PKG (1:1000 (19)), anti-CS (3D11, 1 μg/mL), anti-HSP70 (2E4, 1.0 μg/mL (31)), anti-Exp1 (1:1000, kind gift of Dr. Volker Heussler) and anti-MSP1 (1:400, kind gift of Dr. Anthony Holder). Secondary antibodies (anti-mouse Alexa488, anti-rabbit Alexa594 and anti-chicken Alexa 594, Molecular Probes) were used at a dilution of 1:3000.

Sporozoites (0.5-1 x 10^6^) were purified as previously described(32), air-dried at room temperature on poly L-lysine coated glass slides. They were fixed in 4% PFA for 20 min at RT, permeabilized with 0.5% TritonX-100 for 15min at RT before blocking with 3% BSA in PBS for 1h. Primary antibodies diluted in blocking solution were added at the appropriate dilutions and incubated either for 1h at RT or overnight for 4°C. Secondary antibodies diluted in blocking solution were incubated for 1h at RT. Washes with PBS were performed after incubation with each antibody.

Protein expression in liver stages was examined by infecting HepG2 cells, at 70% confluency, with sporozoites (2×10^4^ – 4 ×10^4^). At the appropriate time post-infection, cells were fixed in 4% PFA for 20 min, permeabilized with cold methanol for 15min and blocked in 3% BSA/ PBS for 1 h. For MSP1 staining, cells were permeablized with 0.1% TritonX-100. Primary antibodies diluted in blocking solution were added at the appropriate dilutions and incubated either for 1 h at RT or overnight for 4°C. Cells were washed 3 times with PBS. Secondary antibodies diluted in blocking solution were added at the appropriate dilutions and incubated for 1 h at RT.

Protein expression in merosomes was examined by infecting HepG2 cells, at 70% confluency, with sporozoites (8×10^4^ – 10 ×10^4^). Medium was replaced every 12h. Detached cell-containing cell culture supernatant was collected at 65 – 67h p.i. and treated with either IAA (500 μM) or vehicle (1% ethanol). After 90 min at RT, cells were allowed to settle onto polyL-Lysine coated slides, fixed in 4% PFA for 20 min and permeabilized with cold methanol for 15 min before treatment with 3% BSA/ PBS for 1 h for blocking. Antibodies were added as described above.

Images were captured using a Nikon A1R laser scanning confocal microscope using 60X/NA1.4 oil objective. Image deconvolution was performed using the Nikon NIS Elements Advanced Research software. For measuring the diameter of liver stages, images were taken at random using a 100x objective on an Olympus BX61 microscope. The area of a region-of-interest was measured using Olympus CellSens software.

### Western blot analysis for protein detection

Sporozoites (0.5×10^6^ CDPK1-aid-HA/ treatment and 5×10^6^ CDPK5-aid-HA/ treatment) were lyzed in Laemlii buffer in the presence of protease inhibitor. Protein lysates were separated on a 12% SDS-PAGE gel and transferred to a PVDF membrane using standard wet transfer. The membrane was blocked in 3% BSA in PBS containing 0.1% Tween20. Anti-HA (Biolegend, 1:1000) and anti-GFP (a kind gift of Dr. Conrad Beckers, 1:1000) antibodies were incubated overnight at 4°C. Secondary antibodies conjugated to Horse Radish Peroxidase (GE HealthSciences) were incubated for 1 h at RT. The membrane was washed (3 ×15 min) after primary and secondary antibodies in PBS containing 0.1% Tween20. Membrane was developed using SuperSignal substrate (Thermofisher Scientific).

### Motility Assays

Sporozoites dissected in DMEM (1×10^4^) were incubated at RT for 90 min with either IAA (500 μM) or vehicle (1% ethanol), in a volume of 25 μL. After addition of equal volume of 6% BSA, they were transferred to a 96-well plate with an optical bottom and centrifuged for 3 min at 4°C. Sporozoites were filmed on a Nikon A1R laserscanning confocal microscope using a 20X/NA0.75 objective at 37°C. Movies were recorded over 90 frames at 1 Hz. Image acquisition and analysis was performed using NIS Elements software from Nikon. Fluorescence intensity projections were processed using NIS Elements and movement patterns were determined through visual inspection of individual sporozoites.

### Cell Traversal Assays

Cell traversal assays were performed as previously described (33). Briefly, sporozoites (4 ×10^4^/well), in plain DMEM medium, were treated with either IAA (500 μM) or vehicle (1% ethanol) in a volume of 150 μL. After 90 min at RT, sporozoites were added to confluent HepG2 cells plated in 8-chamber LabTek slides in the presence of fluorescein-conjugated dextran (1 mg/mL). After incubation at 37°C for 1 h, cells were washed in PBS and fixed with 4% PFA.

### Sporozoite Invasion Assays

Sporozoites (4×10^4^/well), in plain DMEM medium, were treated with either IAA (500 μM) or vehicle (1% ethanol) in a volume of 75 μL. After incubation for 90 min at RT, 75 μL of 2% BSA in DMEM was added along with either IAA (500 μM) or vehicle (1% ethanol). Sporozoites (150 μL/ well) were added to confluent HepG2 cells plated in 8-chamber LabTek slides. Invasion assays were performed as previously described (33). Briefly, 90 min after sporozoite addition, cells were fixed with 4% PFA, blocked with 3% BSA in PBS and incubated with 3D11 (1μg/ mL) for 1h at RT. After washes with PBS, cells were incubated with anti-mouse Alexa594 (1:3000) for 1h at RT. Cells were permeablized with cold methanol for 15 min, blocked and incubated with 3D11. After washes, cells were incubated with anti-mouse Alexa488 (1:3000) for 1h at RT. The number of sporozoites that became intracellular were determined by calculating the difference in numbers of sporozoites that stain with Alexa488 and Alexa594. The percentage of sporozoites that invaded was calculated by determining the percentage of total sporozoites that were intracellular.

### Sporozoite Infection Assays

Sporozoites, pre-treated with IAA (500 μM), vehicle (1% ethanol), 1294 (2 μM) and zaprinast (50 μM), were added to HepG2 cells, for a final volume of 200 μL. Sporozoite-containing medium was replaced with complete DMEM at 2 h p.i. and 24 h p.i. Cells were fixed with 4% PFA at 48 h p.i., permeablized with cold methanol for 15 min, blocked for 1 h with 3% BSA in PBS before incubation with the anti-HSP70 and anti-mouse Alexa488 antibodies as described above. The number of liver stages was determined through microscopic examination.

### Liver stage development

Sporozoites (2 - 4 ×10^4^/well), in DMEM supplemented with 10% FCS, were added to HepG2 cells plated in 8-chamber LabTek slides. To determine its effect on liver stage development, IAA or vehicle was added to infected HepG2 cells 3 h p.i.. Medium was replaced at 14 h p.i. and cells fixed for immunofluorescence analysis using anti-HSP70 as described above.

### Merosome formation

Sporozoites (8 - 10×10^4^/well) were added to sub-confluent HepG2 cells plated on glass coverslips in 24-well plates. Medium was replaced every 12 h. To determine its effect on the development and release of merosomes, IAA was added to infected HepG2 cells 48 h p.i.. The number of merosomes present in the medium was quantified in a hemocytometer at 65 – 67 h p.i..

*In vivo* infections. Merosomes were treated with IAA (500 μM) or vehicle (1% ethanol) for 90 min at RT in DMEM. Following treatment, they were injected intravenously into Swiss-Webster female mice (4-5 mice/group, 6-8 weeks) either immediately (200 intact merosomes/mouse) or after passaging 10 times through a 23G needle (5000 ruptured merosomes/mouse) as previously described (34). Blood was collected from mice 24 h p.i. and analyzed by flow cytometry for the number of GFP-positive events as previously described (35). A total of 1×10^5^ events was counted per sample. Parasitemia was confirmed through daily microscopic examination of Giemsa-stained blood smears.

### Statistical analysis

Data were examined using GraphPad Prism v7.

## Acknowledgments

We thank Dr. Dabbu Kumar Jaijyan and Luke Fritzky for helpful discussion and technical assistance.

---

**Movie 1:** Video of vehicle-treated Ostir1 sporozoites.

**Movie 2:** Video of IAA-treated Ostir1 sporozoites.

**Movie 3:** Video of vehicle-treated CFP-expressing CDPK1-aid-HA sporozoites.

**Movie 4:** Video of IAA-treated CFP-expressing CDPK1-aid-HA sporozoites.

**Movie 5:** Video of vehicle-treated mCherry-expressing CDPK4-aid-HA sporozoites.

**Movie 6:** Video of IAA-treated mCherry-expressing CDPK4-aid-HA sporozoites.

**Movie 7:** Video of vehicle-treated GFP-expressing CDPK5-aid-HA sporozoites.

**Movie 8:** Video of IAA-treated GFP-expressing CDPK5-aid-HA sporozoites.

## Supplementary Information

**Supplementary Figure 1: Characterization of CDPK4-aid-HA and CDPK5-aid-HA parasites. A)** Schematic for tagging with CDPK4 and CDPK5 with aid-HA_2x_ through single recombination. Blue box represents the expression cassette for hDHFR and fluorescent marker. Dotted line represents the site of plasmid linearization. **B)** Integration of targeting constructs was detected by interrogating the locus by PCR. Primer pairs P1 + P2 and P3+P4 were used to detect 5’ and 3’ integration events in CDPK4-aid-HA parasites, respectively. Primer pairs P5 + P2 and P3+P6 were used to detect 5’ and 3’ integration events in CDPK5-aid-HA parasites, respectively. **C)** Sporozoite numbers in salivary glands of mosquitoes infected with CDPK1-aid-HA, CDPK4-aid-HA, CDPK5-aid-HA and isogenic control parasites (Ostir1) used as recipients of the targeting plasmids. The experiment was repeated 3-5 times. **D)** Representative images of CDPK5 expression in CDPK5-aid-HA sporozoites after IAA treatment. IAA efficiently depletes CDPK5 protein in sporozoites. **E)** Cell traversal by sporozoites in unaffected by depletion of CDPK1 or CDPK5. The number of cells containing dextran-FITC (± SEM) formed in each condition was normalized to vehicle-treated controls. The experiment was performed 3-4 times with 4 technical replicates. Results shown are average of 3-4 experiments with 3 technical replicates (± SEM), normalized to vehicle-treated samples.

**Supplementary Figure 2:** Detection of CDPK1 protein in CDPK1 cKO sporozoites. CDPK1 cKO sporozoites were generated through FlpL-mediated deletion of the CDPK1 ORF in FlpL-expressing parasites (FlpL/TRAP). PbCDPK1 was detected using an anti-PfCDPK1 antibody. Images shown are representative of CDPK1 expression in these sporozoites.

**Supplementary Figure 3:** Effect of CDPK4 inhibition and CDPK5 depletion on parasite exit from hepatocytes. **A)** Off-target inhibition of PbPKG by 1294 in sporozoites was interrogated using parasites expressing either HA-tagged wild-type PKG (PKG-HA) or its gatekeeper mutant (T_619_Q-HA). Liver stages were quantified at 48 h p.i. The number of liver stages (± SD) formed in each condition was normalized to vehicle-treated controls. The experiment was performed once with 4 technical replicates. **B)** Dose-dependent inhibition of merosome formation by 1294. Merosomes and detached cells were quantified at 65-68h p.i. with PbLuc sporozoites. Compound was added to infected HepG2 cultures at 48 h p.i. and refreshed every 12 h. The number of merosomes/detached cells (± SEM) formed in each condition was normalized to vehicle-treated controls. The experiment was performed twice with technical triplicates. **C)** Growth rate of CDPK5-aid-HA parasites in erythrocytes. Parasitemias (± SD) of mice infected with vehicle- or IAA-treated CDPK5-aid-HA merosomes were determined daily. Data are from a representative experiment (5 mice/group). Data were analyzed using an unpaired *t*-test, non-significant (n.s.) *P* > 0.05, * *P* value < 0.05.

